# Noninvasive genetic sampling of two flying fox species provides a high rate of genotyping success and a low error rate during amplification

**DOI:** 10.1101/2021.06.15.447175

**Authors:** Mohamed Thani Ibouroi, Ali Cheha, Aurelien Besnard

**Affiliations:** EPHE, PSL Research University, CNRS, UM, SupAgro, IRD, INRA, UMR 5175 CEFE, F-34293 Montpellier, France

**Keywords:** Comoros Islands, frugivorous bats, faecal samples, genetic data, cytochrome b, microsatellites

## Abstract

Noninvasive genetic sampling techniques are useful tools for providing genetic data that are crucially needed for determining suitable conservation actions. Yet these methods may be highly unreliable in certain situations for instance, when working with faecal samples of frugivorous species in tropical areas. In this study, we tested the applicability of noninvasive genetic sampling on two Comoro Islands flying fox species: *Pteropus livingstonii* and *P.seychellensis comorensis* in order to optimize the sampling and laboratory process. Both mitochondrial (mtDNA) and microsatellite markers were tested using two common faeces conservation protocols (ethanol and silica gel), and the polymerase chain reaction (PCR) success and genotyping error rates were assessed. The average proportion of mtDNA PCRs positive results was 55% for *P.livingstonii* and 38% for *P.s.comorensis*, and higher amplification success was obtained for samples preserved in ethanol as compared to silica gel. The average genotyping success rate was high (74% for *P.livingstonii* and 95% for *P.s.comorensis*) and the genotyping error rate was low for both species. Despite our results confirm the effectiveness of using noninvasive genetic sampling methods to study flying fox species, the protocol we used can be optimized to provide higher efficiency. Some recommendations related to field sampling protocols and laboratory methods are proposed in order to optimize amplification rate and minimize genotyping errors.

## INTRODUCTION

Establishing effective habitat management and wildlife conservation measures requires information such as population size, genetic diversity and long-term demographic data (Sharma *et al.*, 2012). Such information has traditionally been gathered using capture–recapture surveys or genetic sampling of tissue or blood, approaches that require the physical capture of individuals, which is time-consuming and difficult for elusive species (Marucco *et al.*, 2009). Moreover, physical capture often disturbs and can sometimes even injure or kill animals, thus posing ethical issues, especially as regards threatened animals. Today, genetic data can be successfully obtained noninvasively from diverse animal sources, such as hair or faecal materials, minimizing the risk and stress for animals (Taberlet *et al.*, 1999). Noninvasive genetic sampling (NIGS) thus represents an alternative option to traditional methods for acquiring genetic data (Marucco *et al.*, 2009). NIGS is easy to implement and is now widely used to assess management and conservation issues for many species, such as brown bears or *Ursus arctos* (Taberlet *et al.*, 1997), insectivorous bats (Puechmaille and Petit, 2007), or large primates (*Pan troglodytes*, Morin *et al.*, 2001). Yet while NIGS has become increasingly popular for wildlife monitoring over the two last decades (Boston *et al.*, 2012), few studies have used these methods to focus on bat populations from tropical and subtropical regions (Baldwin *et al.*, 2010; Ibouroi et al., 2018a). This paucity is likely due to difficulties related to the low quantity and poor quality of DNA contained in certain noninvasive samples. In particular, faecal samples from frugivorous animals are often degraded or contain many PCR inhibitors (Ibouroi et al., 2021). On the one hand, faecal samples of frugivorous animals can retain some non-digested fruit remains that could attract insects. The presence of insects in the faecal samples could speed up the DNA degradation and lead to the deposition of different inhibitors (Palomares *et al.*, 2002). On the other hand, faecal from frugivorous animals are often soft in terms of consistency or even liquid. This could accelerate DNA degradation because of the high level of humidity in the sample (Morin *et al.*, 2001; Palomares *et al.*, 2002). All together, these factors can lead to genotyping errors such as a high rate of allelic dropout (ADO) and false alleles (FA) (Morin *et al.*, 2001). This can result in potential bias in genetic and demographic results (Boston *et al.*, 2012). It is thus crucial to test and improve methods for obtaining genetic data on elusive fruit bat species in tropical regions.

*Pteropus livingstonii* and *P.seychellensis comorensis* are two fruit bat species endemic to the Comoros Islands, where their importance to ecosystem functioning as pollinators and seed dispersers has been demonstrated (Ibouroi et al., 2018b). Despite their key role in maintaining of ecosystem regulation in the Comoros, to date few studies have been undertaken in these islands regarding these species’ genetic diversity or population dynamics, making it difficult to recommend appropriate management plans (Ibouroi et al., 2018a; Ibouroi et al., 2021). This lack of knowledge is partly due to the difficulty of carrying out capture–recapture surveys in these species’ habitats, since individuals are often located on large, tall trees (*P.s.comorensis*, (Ibouroi et al., 2018b) or in highly inaccessible, dense forests (*P.livingstonii*, Ibouroi et al., 2018b; Ibouroi et al., 2021). Bearing in mind the problems related to potential genotyping errors using noninvasive sampling of faeces, we felt this method could be considered as a possible alternative to direct monitoring in order to provide genetic data for the two species, which would then allow the development of relevant conservation measures.

To this end, we tested the applicability of NIGS on these two flying fox species with the following aims: (1) to test the impact of two common faeces conservation protocols (96° ethanol and silica gel), of storing duration and on sampling period and locations on PCR success with mitochondrial DNA cytochrome b; (2) to assess the genotyping success rate and the proportion of genotyping errors (ADO and FA) for both species using nine microsatellite loci; and (3) to optimize the use of such techniques on fruit bat species, especially in tropical regions.

## MATERIALS AND METHODS

Faeces samples were collected at 18 sites (12 sites for *P.livingstonii* and 6 sites for *P.s.comorensis)* of the Comoros archipelago (Anjouan, Grande Comore and Mohéli). At each site, a plastic trap was placed below each roosting tree from 6:00 pm to 8:00 am, collecting up to 15 samples of faecal material. We define faecal sample here as one dropping from an individual of flying fox. These faecal samples were then placed into 5-ml tubes containing different preservation agents: 96° ethanol or silica gel. The samples were kept at room temperature (between 20 and 30 °C) in the Comoros Islands before being transported to France and frozen (−20 °C) until processing for DNA extraction.

The DNA from the samples was extracted at a laboratory facility specializing in the analysis of degraded and sensitive DNA (“ADN dégradé” platform of the LaBex CeMEB, Montpellier, France). The DNA was extracted using a QIAGEN kit (DNeasy mericon Food Kit 69514) following the manufacturer’s protocol. It was amplified through polymerase chain reaction (PCR) using both mitochondrial loci (mtDNA, 460 bp cytochrome b) and 9 microsatellite loci. For the mtDNA amplification, a 10-μL mix containing 1X of the PCR buffer from the ‘QIAGEN Master Mix Kit’, 0.5 μM of the primer CBPL1 (5’ TAC ATC CCA GCC AAC CCA CTA 3’; designed for this study), 0.5μM of the primer H15915 (5’ AAC TGC AGT CAT CTC CGG TTT ACA AGA C 3’; (Irwin *et al.*, 1991)), 3 μl of H2O and 1 μl of DNA was run in a Thermocycler. PCR process followed these conditions: an initial denaturation at 94 °C for 2 minutes, followed by 35 cycles of denaturation (94 °C, 60s), annealing (45 °C, 60s), elongation (72 °C, 90s) and a final elongation at 72 °C for 10mn. PCR products were separated by electrophoresis in 1–2 % agarose gels in TBE buffer. Gels were visualized under UV and photographed with a digital image system. Each sample was examined according to its band from the agarose gel to perform dilution. PCR products were diluted depending on the intensity of each band. For a band of high intensity, we added a volume of 8 μl of H2O plus 2 μl of PCR product to have a final volume of 10 μl of diluted PCR product and for a band of low intensity we added 4 μl of H2O plus all PCR product (~6μl) for sequencing. The diluted PCR products were sent to Eurofins Genomics (Ebersberg, Germany) for sequencing. Electropherograms were visually checked for errors and sequences were assembled using CodonCode Aligner 4.2.5 (CodonCode Corporation). The proportion of PCR success rate was calculated considering the number of samples showing positive sequences divided by the total number of samples. For species identification, we used MEGA software (Tamura *et al.*, 2013) to build a haplotype phylogenetic tree with maximum likelihood methods. A comparison of several models was performed using the Akaike Information Criterion (Akaike 1973) in the program MEGA 6 and the model HKY+G (Hasegawa et al. 1985) was selected as the best model. For the genotyping process, 9 microsatellite loci (described for *P. rodricensis*, a genetically close species to both *P.s.comorensis* and *P.livingstonii*; O’Brien *et al.*, 2007) were tested. All forward primers were fluorescently labelled and combined with reverse primers in three multiplex PCRs at different temperatures (Table 1).

**Table 1:** Properties of the nine multiplexed microsatellite loci: SR.PLV= Allele size range for *P.livingstonii*, SR.PSC= Allele size range for *P.s.comorensis, locus source ; O’Brien et al 2007*

| Multiplex(M) | Locus | Annealing temperature | PRP.CLR | SR.PLV | SR.PSC |
| --- | --- | --- | --- | --- | --- |
| M1 | **A1** | 52 °C | Red | 145–163 | 157–163 |
|  | **C6** | 52 °C | Blue | 208–314 | 283–286 |
|  | **PH9** | 52 °C | Green | 215–265 | 213–241 |
| M2 | **A2** | 50 °C | Blue | 171–187 | 171–189 |
|  | **A8** | 50 °C | Red | NA | NA |
|  | **CSP7** | 50 °C | Green | 250–278 | 258–292 |
| M3 | **A3** | 54 °C | Red | 277–359 | 327–361 |
|  | **B29** | 54 °C | Blue | 210–254 | 224–330 |
|  | **PH4** | 54 °C | Black | NA | NA |

Each PCR mix for each multiplex contained a final volume of 10 μl: 5 μl of PCR buffer ‘2x QIAGEN Multiplex PCR Master Mix, ref. 206145’ (2X master mix, 1.5 μM MgCl 2, Taq, dNTPs), 0.2 μM of the primer mixes, 1 μl of DNA and 3 μl of H2O. The PCR mix was also run in a thermocycler Eppendorf (Mastercycler epgradient S) in the following conditions: an initial denaturation at 95 °C for 15 min, followed by 35 cycles of denaturation (94 °C, 30 s), annealing (90 s at 52 °C, 50 °C and 54 °C respectively for multiplex 1, multiplex 2 and multiplex 3, Table s3), elongation (1 min at 72 °C), and a final elongation step of 30 min at 60 °C. Products were run in a 16 capillary sequencer (3130 xl Genetic Analyzer, Applied Biosystems) at the platform “Genotypage-Séquençage” of the LabEx CeMEB with the LIZ500standard size (omitting the 250bp peak) and the alleles of each sample were scored with GeneMapper v4.5 (Applied Biosystems). To assess the reliability and error rates of the genotyping of the faecal samples, each sample was independently genotyped using the multiple-tubes procedure (Taberlet *et al., 1997*). PCR were run at least three times. In some cases no amplification occurred for at least one of the three runs. In this situation, we carried out some additional runs for the corresponding samples to obtain three or more positive PCR. The genotyping error rate was assessed with the GIMLET v1.3.3 program (Valière, 2002) and was classified as (1) allelic dropout (ADO) when heterozygous alleles from at least three independent repeated genotyping procedures showed homozygous alleles, or (2) false alleles (FA) if a homozygote from a triplicate genotyped set showed heterozygote alleles (Valière *et al.*, 2003). Consensus genotypes were assessed using the GIMLET program.

### Effect of the conservation method, sampling period and preservation time on the mtDNA success rate

During our field sampling, we did not get the same sample size for ethanol and silica gel. The main objective of our field session was to collect reliable samples for genotyping and for microsatellite analysis. After the first field session, the preliminary results of some amplification tests showed that ethanol was more efficient in terms of sample conservation than silica gel, then during our last field session, all samples were stored in ethanol only. To assess the effect of preservation time, sampling period, Island where faecal were collected and the conservation protocol on the mtDNA PCR success rate, we performed a Generalized Linear Model with binomial distribution and logit link. As we did not have sufficient number of samples for faeces preserved in silica gel during the second and third sampling sessions, tests were performed only on samples stored in ethanol for preservation time, Island effect and sampling period. As silica gel samples were all extracted 3 to 4 months after collection, we tested for the effect of preservation method on samples stored 12, 13 and 14 weeks in both ethanol and silica gel. For the effect of sampling period, samples collected between November to February (high rainfall season) were compared to samples collected between March to September (dry season). All statistic tests were performed for *P.livingstonii* only as we did not have sufficient samples to perform statistical analyses on *P.s.comorensis*. Test was considered significant if p-value<0.05. All statistical analysis were performed using R (R Development Core Team, 2016).

## REUSLTS

### PCR success with mtDNA

A total of 337 faecal samples were collected on the three islands of Comoros: 244 for *P.livingstonii* (178 from Anjouan Island and 66 from Mohéli Island) and 93 samples for *P.s.comorensis* (20 from Anjouan, 50 from Grande Comore and 23 from Mohéli). The average proportion of mtDNA PCRs with positive results was 55% (135/244) for *P.livingstonii* compared to 38% (36/93) for *P.s.comorensis* (Table 2).

**Table 2:** MtDNA amplification success rate using different DNA preservation methods (top of table) and microsatellite genotyping success per locus (bottom of table): ADO = allelic dropout; FA = false alleles rate; GTS = frequency of genotyping success; GTSR = percentage of genotyping success, NA = not available

| mtDNA |  | *P.livingstonii* |  |  | *P.s.comorensis* |  |  |  |
| --- | --- | --- | --- | --- | --- | --- | --- | --- |
|  | Ethanol 96° | Silica gel | Total |  | Ethanol 96° | Silica gel | Total |  |
| Anjouan | 74/115<br>(64%) | 19/63<br>(30%) | 93/178<br>(52%) |  | 13/20<br>(65%) | NA | 13/20<br>(65%) |  |
| Mohéli | 25/32<br>(78%) | 17/34<br>(50%) | 42/66<br>(63%) |  | 1/5<br>(20%) | 6/18<br>(33%) | 7/23<br>(33%) |  |
| Grande Comore | NA | NA | NA |  | NA | 16/50 (32%) | 16/50<br>(32%) |  |
| Total | 99/147<br>(67%) | 36/97<br>(37%) | 135/244<br>(55%) |  | 14/25<br>(56%) | 22/68<br>(32%) | 36/93<br>(38%) |  |
| Microsatellites |  | *P.livingstonii* |  |  | *P.s.comorensis* |  |  |  |
| Locus | GTS | GTSR | ADO | FA | GTS | GTSR | ADO | FA |
| A1 | 90/120 | 75% | 0.014 | 0.000 | 21/22 | 95% | 0.000 | 0.000 |
| C6 | 89/120 | 74% | 0.012 | 0.000 | 22/22 | 100% | 0.000 | 0.000 |
| PH9 | 87/120 | 73% | 0.042 | 0.000 | 22/22 | 100% | 0.000 | 0.000 |
| A2 | 90/120 | 75% | 0.005 | 0.027 | 22/22 | 100% | 0.000 | 0.000 |
| CSP7 | 89/120 | 74% | 0.039 | 0.010 | 22/22 | 100% | 0.028 | 0.010 |
| A3 | 88/120 | 73% | 0.016 | 0.020 | 16/22 | 72% | 0.00 | 0.038 |
| B29 | 90/120 | 75% | 0.000 | 0.015 | 22/22 | 100% | 0.00 | 0.000 |
| Total | 89/120 | 74% | 0.018 | 0.010 | 21/22 | 95% | 0.004 | 0.020 |

Statistical analyses revealed that conservation protocol had a strong influence on the mtDNA success. mtDNA PCR amplification success was significantly lower in silica-gel (0.37 95%CI [0.28-0.47]) than in alcohol (0.79 95%CI [0.59-0.91], on samples stored for 12, 13 and 14 weeks, z=14.22, df=1, p<0.001). Storing duration have no significant impact on the DNA preservation in Ethanol (z= 0.16, ddf =1 p-value=0.68, the amplification rate varied from 0.70 for 4 weeks to 0.64 for 14 weeks). According to our statically analysis, the sampling period significantly impacted mtDNA success rate, mtDNA success rate was significantly lower for samples collected during the wet period (0.58, 95%CI [0.49-0.68]) compared to dry period (0.84 95%CI [0.71-0.91], z= 8.12, df=3 p-value=0.04). When mtDNA amplification success was modelled depending on the Islands, mtDNA success rates significantly differed between Anjouan (0.64 95%CI[0.55-0.72]) and Moheli (0.80 95%CI [0.62-0.91], z= 8.12, df =3 p-value=0.04).

### Phylogenetic analysis with mtDNA

We built a phylogenetic tree with 156 amplified sequences obtained from faeces (resulting in a total of four haplotypes for *P.s.comorensis* and 11 haplotypes for *P.livingstonii*) as well as 20 cytochrome b sequences from GenBank (10 sequences for *P.livingstonii*, and 10 sequences for *P.s.comorensis*). The resulting tree (Haplotype phylogenetic tree as well as all GenBank sequences are shown in electronic supplementary material) confirmed that all samples belonged to one of the two species, but also revealed that eight samples collected under *P.livingstonii* dormitory trees and assigned to this species in the field were in fact from *P.s.comorensis* (identification error of 5%). All sequences are available in GenBank (accession numbers: MF471481 - MF471633).

### Genotyping success and error rates with microsatellite loci

To increase the probability of positive PCR results and to minimize costs and time, only the samples with positive results for mtDNA PCR were genotyped. This allowed us to ensure that we amplified faecal samples with sufficiently non-degraded DNA.

From the 171 positive mtDNA samples (for the two species), microsatellites were amplified for 138 of these, including 120 out of 135 positive mtDNA samples for *P.livingstonii*, and 22 out of 36 positive mtDNA samples for *P.s.comorensis*.

Genotype consensus was established with 7 loci (Table 2) on 90 out of 120 samples for *P.livingstonii* compared to 21 out of 22 samples for *P.s.comorensis*. The two remaining loci (A8 and PH4) did not provide reliable genotypes and were discarded in further analysis. The genotyping success rate was high for all loci for both species, varying from 72% (locus A3 and PH9) to 75% (locus A1, A2 and B29) with an average of 74% for *P.livingstonii*, and from 72% (A3) to 100% (C6, PH9, A2, CSP7 and B29) with an average genotyping success rate of 95% for *P.s.comorensis*. The overall genotyping error rate was low for both species. The average proportion of allelic dropout (ADO) and false alleles (FA) appeared to vary between markers (Table 2). For *P.livingstonii*, the frequency of ADO and FA were respectively 1.8% and 0.1%. An average of 0.4% and 2% of ADO and FA respectively was recorded for *P.s.comorensis* (Table 2).

## DISCUSSION

Results presented (Table 2) indicate moderate and low mtDNA PCR success rate for *P.livingstonii* and *P.s.comorensis* respectively. These results are however comparable to those obtained on certain mammal species using mtDNA, including insectivorous bats in Europe (*Myotis capaccinii*: 50%, Viglino *et al.*, 2016), but are lower than those obtained for others (*Myotis mystacinus* and *Myotis nattereri:* 72–88%, Boston *et al.*, 2012). These contrasting results could be explained by differences in diet of the studied species. According to many studies, the PCR success rate is highly affected by different PCR inhibitors that can occur in faecal materials depending on the diet of the species, and these inhibitors are probably in high quantity in faeces of frugivorous animals (Puechmaille and Petit, 2007, Morin et al 2001). Then the type of habitat where the samples were collected and the sampling conditions can impact DNA preservation. Some studies indicate for instance that roost conditions (e.g. humidity, temperature, etc.) as well as abundances of insects and rainfall can impact DNA preservation (Boston *et al.*, 2012; Viglino *et al.*, 2016). A last, the interval between the time the samples were collected and the DNA were extracted, and/or the protocol used to preserve the sample (ethanol or silica gel) and the conditions of DNA extraction (Boston *et al.*, 2012) can also strongly impact amplification success.

### Impact of sampling condition, preservation methods and storing duration on amplification success rate

According to previous studies (Murphy *et al.*, 2003), high humidity could have an effect on DNA conservation in faecal samples leading to low PCR success rate and it might explain the relatively low mtDNA amplification success we obtained. It is difficult to provide a precise value of the humidity in the forest during data collection since we did not collect this information during field sessions. Yet according to the Climatetemp.com, the studied Islands have an average humidity of 76% during wet season. This average value at the island scale probably represents the lowest value we get during the field since several samples (especially those of *P. livingstonii*) were collected in rainy forest at high altitude. Moreover, during the sample collection, it rained almost every night and the weather was fogy and very humid. We tested the relationship between the mtDNA success rate and the Island where samples were collected and our results highlighted a low but significant difference between the islands. Samples collected in the forests of Anjouan showed a lower rate of mtDNA amplification than those of Moheli. The forests of Anjouan are notably characterized by higher rainfalls and humidity than forests of the Island of Moheli which are characterized by moderate rainfalls (UNEP 2002). The impact of humidity is also supported by our analysis regarding sampling period. Samples collected during dry periods show higher mtDNA amplification success than samples collected during rainfall periods. This is probably due to the fact that rainfall periods are characterized by high humidity; of course; but also high temperatures, and a high abundance of insect that could impact DNA (Palomares *et al.*, 2002).

Our results show higher mtDNA amplification success for samples preserved in ethanol as compared to silica gel. These results are in line with those obtained by Roeder *et al*. (2004) showing that faecal samples in silica gel were poorly preserved for tropical species. In contrast, Puechmaille and Petit (2007) obtained high amplification success rates for samples preserved in silica gel for insectivorous bats in Europe (Goodman *et al.*, 2012; Puechmaille and Petit, 2007; Ruedi *et al.*, 2012). In our case, the low mtDNA amplification success rate from samples stored in silica gel in comparison to those stored in ethanol can also be explained by the very high level of humidity of the collected samples. Because our samples were exposed to a high level of humidity and mildews, silica gel probably desiccates faeces too slowly and may have not been able to prevent further DNA degradation, as opposed to ethanol (Murphy *et al*., 2002).

According to Murphy *et al.* (2002), faecal samples that have been preserved for a long time (up to six months) show lower success rates for mtDNA amplification (Brinkman *et al.*, 2009). Yet Puechmaille *et al.* (2007) found no difference in mtDNA amplification success between samples stored 6 month versus 18 month in silica gel prior to DNA extraction (Puechmaille *et al.*, 2007). In our case, due to administrative procedures related to exporting the samples, samples were stored at ambient temperature for a long period (between 3 and 4 months) before being processed in the lab. However, our statistical analysis testing for the impact of storing duration on amplification success was not significant as in Puechmaille *et al.* (2007), on shorter period though.

### Effect of laboratory process on mtDNA success rate

Low mtDNA success rate can also be explained by the DNA extraction protocol. We used the DNeasy Mericon Food with our faecal samples as this Kit had already been successfully tested with mammal faeces samples in our lab. Testing different extraction kits and performing some modifications of the extraction protocol are likely to improve the rate of successful extraction as shown in many other species. Hence testing different extraction protocols is recommended to select the most efficient one for the target species (e.g. Puechmaille *et al.* 2007).

Amplifying a shorter mtDNA fragment could also have resulted in a higher amplification success (Broquet *et al.*, 2007). Yet our microsatellite loci ranged from 170 to 345 bp (see O’Brien *et al.*, 2007), we thus assumed that a successful amplification of 400-450bp of mtDNA, should be a relevant proxy of amplification success of these microsatellites. When working with shorter microsatellite loci or if the goal of the study is to test different protocols with mtDNA, we recommend the use of shorter sequences fragment that could lead to higher amplification success and less DNA extract being discarded before genotyping.

### Importance of testing samples with mtDNA before genotyping

Our phylogenetic results highlights that the two species can share the same roosting trees even in sites at which *P.livingstonii* is believed to live alone. This finding highlights the importance of testing samples with mtDNA before genotyping when working with NIGS. Testing with mtDNA amplification not only allows the identification of the species to be confirmed, but also enables the selection of the samples with the best potential DNA for subsequent microsatellite amplifications. These steps could in turn allow reducing genotyping errors, the main challenge in noninvasive genetic sampling. However, identifying species using mtDNA analysis is possible only when the species can be diagnosed with mtDNA markers (Afonso *et al.*, 2017; Puechmaille and Teeling, 2014). This is not the case when introgression occurs on the targeted species that can conduct to misidentification of the species (Puechmaille and Teeling, 2014). In our case, since microsatellite loci analyses were expensive compared to mtDNA, the use of the mtDNA was the best option to verify the presence of DNA in faecal samples and to identify species despite the extra time needed for this step. Indeed, studies of O’Brien *et al.* (2009), Chen *et al.* (2011) and Almeida et al. (2014) showed that mtDNA allows successfully identifying the two targeted species using cytochrome b gene (Almeida *et al.*, 2014; Chan *et al.*, 2011; O’Brien *et al.*, 2009). Yet, in some situations (when the use of mtDNA is not relevant to separate some species or when working on a single species) the use of microsatellites can be sufficient to select for good quality DNA (Afonso *et al.*, 2017).

### Genotyping and error rates with microsatellite loci

We successfully amplified only 7 microsatellites loci among the 9 tested (Table 1). Two microsatellites markers (A8 and PH4) did not amplify well probably because of some inadequate PCR conditions or non-specificity in the target species.

Our genotyping success rates (Table 2) for both species are high in comparison to those obtained by Baldwin *et al.*, (2010) for the grey-headed fox (*Pteropus poliocephalus*: 57%) and by Viglino *et al.*, (2016) for mouse-eared bats (*Myotis capaccinii, M. emarginatus* and *M. daubentonii*: 53%). However, our results are comparable to those obtained in other studies for different mammal species: European insectivorous bats (*Rhinolophus hipposideros:* 96%, Puechmaille and Petit, 2007; *Myotis mystacinus:* 72–88% and *Myotis nattereri:* 72–90%, Boston *et al.*, 2012) and large primates (*Pan troglodytes:* 79%, Morin *et al.*, 2001).

Two main hypotheses could explain the high genotyping success rate obtained in our study. First, using only mtDNA-positive samples for microsatellite genotyping allowed good quality samples to be selected prior to genotyping, increasing the success rate. Second, the high rate of genotyping success and low genotyping error rate obtained during our study (table 2) could be explained by DNA extraction protocols and conditions, which are, according to Boston *et al.* (2012), two of the most important factors impacting genotyping success. Furthermore, the DNA was extracted using a specialized kit for food DNA extraction, which has been recommended for noninvasive DNA extraction (Zarzoso-Lacoste *et al.*, 2013). Using an appropriate extraction kit allows genotyping errors to be reduced, which increases the accuracy of genetic results (Puechmaille and Petit, 2007; Boston *et al.*, 2012).

## CONCLUSION

Our results confirm the potential of using NIGS to provide genetic and demographic data on frugivorous tropical bats, including flying foxes; data which is urgently needed in order to recommend relevant conservation actions. The risk of NIGS is that an accumulation of genotyping errors can lead to bias and false interpretations with potentially negative consequences. For example, a high rate of false alleles might lead to inflated observed heterozygosity, incorrect genetic diversity or inaccurate population size estimations, resulting in inappropriate conservation management measures. It is thus crucial to develop relevant protocols for such noninvasive sampling techniques. Despite showing that NIGS can be applied to flying foxes in the wild, our work focused on feasibility rather than optimality. Indeed, other studies have demonstrated that different DNA extraction protocols can perform very differently in different situations/with different species (Puechmaille et al. 2007) hence different extraction Kits and preservation protocols should be tested and optimised to increase amplification rate.

Collecting faecal sample from frugivorous animals such as flying fox in humid forest during the wet season is challenging because faeces are relatively liquid due to the frugivorous diet and because of the rainfall. In such conditions, modifications to our collection protocol, by using a cloth just above the plastic to allow faecal samples to dry quickly by filtration, are recommended. Sampling during the dry season is recommended when possible. When using silica gel conservation method, samples should be checked regularly to verify for potential saturation and silica-gel replaced if necessary.

According to Taberlet *et al*. (1997), the use of a multiple-tubes approach allows the genotyping error rate to be verified, increasing result accuracy and minimizing false interpretations. However, this protocol can be time-consuming and costly if a pilot study is not carried out before genotyping. We recommend the use of mtDNA amplification to first identify samples with good quality DNA for the targeted species if species can be identified with mtDNA and to select these before genotyping, thus reducing subsequent time and costs.

## Acknowledgments

We would like to thank the Comoros National Direction of Environment and Forests, the Earth Sciences Department of the University of the Comoros for permission to carry out fieldworks and to export faeces samples (authorization number 002/KM/15/DNEF). Much of the fieldwork was funded by the Rufford Foundation through a Research Support Grant (grant nos. 26731-2 for M.T.I. and 19010-1 for A.C.). Lab analysis are partly funded by the Islamic Development Bank (IDB). The data used in this study was generated at the facility of the platform of “Genotypage-Séquençage” and “ADN dégradé” of the LabEx CeMEB Montpellier, France.

